# Tamoxifen transiently disrupts estrous cyclicity without altering long-term ovarian aging trajectories

**DOI:** 10.64898/2026.09.03.749251

**Authors:** Sarai Badillo, Ewa Poljanska, Sarah Ray, Hana Kilani, Jillian E.J. Cox, Sunghwan Ko, Subhasri Biswas, Doris M. Benbrook, Michael B. Stout, Sarah R. Ocañas

## Abstract

The ovary is both one of the earliest organs to functionally age in the body, with declines emerging well before reproductive failure and contributing to systemic aging. Since ovarian function depends on tightly regulated hormonal and inflammatory cycles, even subtle disruptions can confound aging-related phenotypes. Tamoxifen-inducible Cre systems are widely used to study ovarian biology, however, tamoxifen is a selective estrogen receptor modulator capable of disturbing ovarian physiology. This introduces a critical and often overlooked concern that tamoxifen-based models may produce lasting effects that obscure true biological signals, particularly in ovarian aging studies. To address this, we tested the hypothesis that tamoxifen induces transient physiological disruption without altering long-term transcriptional outcomes. Female mice were treated with tamoxifen or vehicle at 3 months of age, and estrous cyclicity was monitored longitudinally by vaginal cytology at 3 days, 2 months, 3 months, 6 months, and 12 months post-induction. Ovaries were collected at young (6 months) and aged (12 months) time points for bulk RNA sequencing, followed by differential gene expression and pathway-level analyses. We found that tamoxifen treatment disrupted estrous cyclicity shortly after administration, confirming short term physiological effects. However, normal cycling was restored by 3 months post-treatment, indicating recovery of ovarian function. At the transcriptional level, minimal differences were observed between tamoxifen and vehicle treated groups at both young and aged time points. In contrast, aging-associated transcriptional programs were conserved across treatment conditions with shared alterations in pathways related to extracellular matrix remodeling, senescence, and cellular homeostasis. Together these findings demonstrated that while tamoxifen disturbs ovarian physiology in the short term, it does not produce lasting transcriptional effects or aging associated phenotypes. These results resolve a critical methodological concern and provide validation for the use of tamoxifen-inducible systems in ovarian research, supporting their application in studies of reproductive aging and systemic female health when proper controls are integrated into study design.

## INTRODUCTION

The ovary is among the earliest organs in the body to age [1] with functional decline occurring well before complete reproductive failure thus contributing to broader systemic aging and chronic disease development [2, 3]. Consequently, understanding the mechanisms that drive ovarian aging is essential not only for improving women’s reproductive health, but also for defining how reproductive aging influences overall health span. Ovarian aging extends beyond progressive depletion of the follicular reserve and is accompanied by alterations in hormonal signaling, stromal remodeling, fibrosis, chronic inflammation, and the accumulation of multinucleated giant cells (MNGCs) [4]. Importantly, many of these age-associated changes occur within the ovary, which undergoes continual cyclical remodeling throughout the reproductive lifespan [5, 6]. As a result, experimental perturbations that interact with hormonal or inflammatory signaling may have transient effects on ovarian physiology, making it essential to determine whether these changes resolve or persist in ways that could influence the interpretation of aging studies.

Tamoxifen (TAM)-inducible Cre/ERT2 systems are widely used to study gene function in basic and translational research because they enable temporally controlled recombination in adult tissues, which reduces confounding effects from developmental gene manipulation[7, 8]. In these systems, TAM activates Cre-mediated recombination by binding to a modified estrogen receptor ligand-binding domain [9]. However, TAM is also a selective estrogen receptor modulator (SERM) and can alter estrogen signaling in a tissue-specific manner, independent of its role in recombination [10]. This is particularly relevant in estrogen-responsive tissues, including the ovary, where estrogen signaling regulates folliculogenesis, estrous cyclicity, ovulation, and tissue remodeling [11, 12]. Prior studies have shown that although TAM administration can induce transient physiological effects, it does not produce long-lasting alterations in the brain transcriptome or epigenome [13]. However, the ovary may be particularly sensitive to TAM exposure because of its dependence on cyclic estrogen signaling and continual tissue remodeling across the reproductive lifespan [14].

Despite the widespread use of TAM-inducible models in reproductive aging and cancer research [15–21], the short– and long-term consequences of TAM exposure on ovarian physiology and aging outcomes remain poorly defined. This represents an important gap because persistent TAM-induced changes in estrous cyclicity, tissue architecture, or transcriptional state could confound interpretation of studies of ovarian aging, reproductive senescence, or inflammation-associated phenotypes, all of which are associated with ovarian cancer development [22]. Ovarian cancer is considered a disease of aging with mortality increasing sharply after 40 years old [23]. Distinguishing transient TAM effects from lasting alterations is therefore essential for the appropriate design and interpretation of TAM-inducible models in ovarian aging and cancer studies.

In this study, we investigated the effects of TAM administration on estrous cyclicity, ovarian histological aging features, and ovarian transcriptional profiles across aging in female Cx3cr1^Cre/ERT2+/-^NuTRAP^flox/WT^ mice. Our findings show that TAM temporarily disrupts estrous cyclicity; in contrast aging remains the main factor driving changes in ovarian tissue architecture and gene expression independently from TAM exposure. These data support the use of TAM-inducible models in ovarian aging studies when appropriate vehicle controls and sufficient post-TAM recovery periods are incorporated into the experimental design.

## METHODS

### Animals

All animal procedures were approved by the Institutional Animal Care and Use Committee at the Oklahoma Medical Research Foundation and performed in accordance with the National Institutes of Health (NIH) Guide for the Care and Use of Laboratory Animals. Breeder mice were purchased from the Jackson Laboratory (Bar Harbor, ME), bred, and housed at the Oklahoma Medical Research Foundation under SPF conditions in a HEPA barrier environment on a 14/10 light/dark cycle (lights on at 6:00 am). Cx3cr1-cre/ERT2^+/+^ males (Stock #020940) [24] were mated with NuTRAP^flox/flox^ females (Stock #029899) [25] to generate the desired progeny, Cx3cr1-cre/ERT2^+/wt^; NuTRAP^flox/wt^ (Cx3cr1^Cre/ERT2^NuTRAP), as previously performed [15]. The Cx3cr1-NuTRAP model was used as a readily available example of a TAM-inducible Cre system relevant to ovarian aging studies. In this model, TAM-induced recombination enables labeling of tissue-resident macrophages (TRMs) while avoiding long-lasting labeling of circulating monocytes [26], allowing TRMs to be distinguished from monocyte-derived macrophages (MDMs). This inducible approach is important because constitutive macrophage-targeting Cre drivers would label both resident and recruited macrophage populations, limiting the ability to separate TRM– and MDM-associated effects. DNA was extracted from mouse ear punch samples for genotyping. Genotyping was performed using standard PCR detection of Cx3cr1-cre/ERT2 (Jax protocol 27232; primers: 20669, 21058, 21059) and NuTRAP floxed allele (Jax protocol 21509; primers: 21306, 24493, 32625, 32626), as previously described [15]. At time of euthanasia, mice were deeply anesthetized with isoflurane, terminal blood collection was performed by cardiac puncture, and animals were then euthanized by transcardial perfusion with phosphate-buffered saline (PBS) in accordance with AVMA Guidelines for the Euthanasia of Animals.

### Experimental Design

At 3 months of age (mo), female Cx3cr1-NuTRAP mice were randomized to either receive TAM or vehicle treatment and then aged to 6 mo, the young time point or 12 mo, the aged timepoint (n=5-7/treatment/age). Three days following TAM administration, vaginal lavage was performed for twelve consecutive days to assess estrous cyclicity. Estrous cyclicity was also assessed 2, 3, and 9 months post-TAM. At 6 or 12 mo, terminal blood collection was performed, and ovarian tissue was isolated for downstream transcriptomic and histological analyses.

### Tamoxifen

At 3 mo, mice received a single daily intraperitoneal (i.p.) injection of 100 μL TAM in 100% sunflower seed oil for five consecutive days (100 mg/kg body weight, 20 mg/ml stock solution, Sigma, St. Louis, MO), as previously performed [13]. TAM was sonicated in sunflower seed oil prior to injection to ensure proper solubilization. Control mice received 100 μl of 100% sunflower seed oil by i.p. delivery on the same dosing schedule to serve as a vehicle (VHL) control.

### Estrous Cycle Staging

For all mice used in the study, vaginal cytology was performed daily for 12 consecutive days at the same time of day to assess estrous cyclicity at four different time points post-TAM or VHL administration, including: 3 days, and 2, 3, and 9 months following treatment. Vaginal lavage was performed as previously described [27]. Briefly, vaginal cytology was assessed by gentle vaginal lavage with sterile PBS, followed by staining using the Hema 3 Stat Pack (Fisherbrand, Norcross, GA) according to the manufacturer’s instructions. Slides were evaluated under a light microscope to determine estrous cycle stage based on the relative abundance of nucleated epithelial cells, cornified epithelial cells, and leukocytes [28]. Cycle regularity was classified according to the duration spent in each cycle phase, based on previous literature [29]. Mice were considered regular if they spent ≤3 consecutive days in any one stage, irregular if they spent 4-7 consecutive days in any one phase, and acyclic if they remained in a single phase for ≥8 consecutive days.

### Histological analyses

Following cardiac perfusion with ice-cold PBS, one ovary was fixed in 4% PFA in 1X PBS with 1.5% glutaraldehyde for 4 hours, dehydrated through graded ethanol, cleared in xylene, embedded in paraffin, and sectioned for histological analyses. For follicular counting, serial sectioning was performed on the whole tissue to collect one tissue section from every six sequential sections through the whole ovary. Picrosirius Red (PSR) staining for collagen deposition and Sudan Black staining for lipofuscin accumulation was performed on one randomly selected mid-ovarian section per sample and analyzed as previously described [30, 31].

### Plasma 17β-Estradiol Measurement

Plasma 17β-estradiol (E2) concentrations were measured using the DetectX Serum 17β-Estradiol Enzyme Immunoassay Kit (Arbor Assays, KB30-H) based on the manufacturer’s recommendations. Briefly, plasma samples were diluted 1:20 in provided buffer and analyzed in duplicates. A six-point E2 standard curve ranging from 3.75 – 120 pg/mL was prepared according to the manufacturer’s instructions.

Diluted plasma samples or standards (100 µL) were added to the antibody-coated 96-well plate, followed by 25 µL of E2-peroxidase conjugate and 25 µL of E2-specific antibody. Plates were sealed and incubated for 2 h at room temperature with shaking at 800 rpm. Wells were subsequently decanted and washed four times with 300 µL of wash buffer. TMB substrate (100 µL) was added to each well and incubated for 30 min at room temperature without shaking, after which the reaction was terminated by addition of 50 µL stop solution. Absorbance was measured at 450 nm using GloMax Explorer (Promega, 9720100152).

E2 concentrations were determined from the standard curve using a four-parameter logistic (4PL) regression model. Duplicate optical density measurements were averaged, corrected for non-specific binding, and sample concentrations were multiplied by the corresponding dilution factor to obtain the E2 concentration in the original plasma sample. For the analysis, vehicle (VHL)– and tamoxifen (TAM)-treated samples were paired based on their estrous cyclicity.

### RNA Extraction

Following cardiac perfusion with PBS, one ovary was snap-frozen and stored until RNA extraction. To extract RNA, 350 µL of RLT Buffer (Qiagen, Valencia, CA) with β-mercaptoethanol was added to the frozen ovarian tissue and then homogenized by bead mill (TissueLyser II; Qiagen). Following homogenization, 350 µL of 100% ethanol was added to the sample and then RNA was extracted using the RNeasy Mini Kit (#74106, Qiagen). RNA was quantified with a Nanodrop One^C^ spectrophotometer (#ND-ONEC-W, ThermoFisher Scientific) and quality-assessed by HSRNA ScreenTape (#5067-5579, Agilent Technologies) with a 4150 TapeStation analyzer (#G2992AA, Agilent Technologies).

### Library Construction and RNA-Sequencing (RNA-Seq)

Directional RNA-Seq libraries were made according to the manufacturer’s protocol from 50 ng RNA, as previously described [26]. Poly-adenylated RNA was captured using NEBNext Poly(A) mRNA Magnetic Isolation Module (#NEBE7490L, New England Biolabs) and then processed using NEBNext Ultra II Directional Library Prep Kit for Illumina (#NEBE7760L, New England Biolabs) for the creation of cDNA libraries. Library sizing was performed with HSD1000 ScreenTape (#5067-5584, Agilent Technologies) and quantified by Qubit dsDNA HS Assay Kit (#Q32851, ThermoFisher Scientific) on a Qubit 4 Fluorometer (#Q33226, ThermoFisher Scientific). The libraries for each sample were pooled at 4 nM concentration and sequenced using an Illumina NovaSeq X Plus 25B PE150 flow cell (Illumina, San Diego, CA, USA).

### Data Analysis

RNA-seq reads were aligned to the reference genome using STAR [32] with Ensemble gene annotations. Resulting alignment files were sorted and converted to BAM format using Samtools [33]. Gene-level read counts were generated from BAM files using featureCounts [34] to produce a count matrix for downstream analysis. Differential gene expression analysis was performed in DESeq2 using Bioconductor. Counts were normalized within DESeq2 [35], and statistical testing was performed to identify differentially expressed genes. P-values were adjusted for multiple comparisons using the Benjamini-Hochberg false discovery rate method. Heatmaps were made using Morpheus (https://software.broadinstitute.org/morpheus/) and venn diagrams were made using Venny (https://bioinfogp.cnb.csic.es/tools/venny/) and BioVenn (https://biovenn.nl/). Lists of differentially expressed genes were imported into Ingenuity Pathway Analysis (IPA) 01.12 (Qiagen Bioinformatics) to assess upstream regulator, canonical pathway, as well as diseases and functions analyses. Volcano plots were generated with the R package EnhancedVolcano using the differentially expressed genes as the input. The PCA was performed in R with the prcomp and visualization was done with the autoplot function from the R package ggfortify. The boxplots were made with ggplot in R. The scatter plot was made with the lm function and viewed with geom_point based off log10 normalized data.

### Statistics

Datasets were analyzed using GraphPad Prism version 11.0 and represented as dot plots with underlying bar graphs, line graphs, bar graphs, or box graphs with mean ± S.E.M (standard error of the mean). All statistical tests used in data analysis are specified in their respective figure. Generally, discrete data were assessed for normality, followed by an appropriate parametric or nonparametric test selected based on the number of groups being compared. p<0.05 was considered statistically significant.

### Data and Code Availability

Raw and processed RNA-seq data generated in this study are deposited in NCBI GEO under accession number GSEXXXX. Analysis code used to generate the figures will be made available through a public GitHub repository or equivalent data archive upon publication. Additional data supporting the findings of this study are available from the corresponding author upon reasonable request.

## RESULTS

Tamoxifen (TAM) is widely used to induce recombination in Cre/ERT2-based mouse models. However, because TAM functions as a selective estrogen receptor modulator (SERM), it may exert tissue-specific effects on estrogen signaling. This is particularly relevant in the ovary, where estrogen signaling is essential for estrous cyclicity and ovarian function. The goal of this study was to determine whether TAM administration has long-lasting effects on estrous cyclicity or ovarian aging outcomes that could confound interpretation of TAM-inducible models in studies of ovarian aging.

### TAM administration induces temporary disruption in the murine estrous cycle

Female Cx3cr1^Cre/ERT2^NuTRAP mice were treated with TAM or VHL at 3 mo and periodically monitored for estrous cyclicity until 12 mo. Ovarian tissue was collected at young (6 mo) and aged (12 mo) to assess the effects of TAM on transcriptomic and histological outcomes (**Fig. 1A**). Estrous monitoring revealed a transient disruption of cyclicity following TAM administration (**Fig. 1B-I**). This disruption was most evident immediately after TAM treatment (**Fig. 1B,F**). Specifically, TAM-treated mice remained predominantly in metestrus during the direct post-treatment monitoring period (**Fig. 1B**), resulting in a complete cessation of cycling across the 12-day observation window (**Fig. 1F**). By 5 mo (approximately 2 months after TAM treatment) most TAM-treated mice had resumed cycling, although a subset exhibited prolonged estrus and irregular cyclicity (**Fig. 1C,G**). By 6 mo (approximately 3 months after TAM treatment) estrous cyclicity was indistinguishable between TAM– and VHL-treated mice (**Fig. 1D,H**). By 12 mo, there remained no difference between TAM– and VHL-treated mice in the proportion of time mice spent in each cycle phase (**Fig. 1E**). However, a subset of mice in both treatment groups exhibited irregular cycling **(Fig. 1I)**, consistent with expected age-associated changes in cyclicity [36].

**Figure 1.**
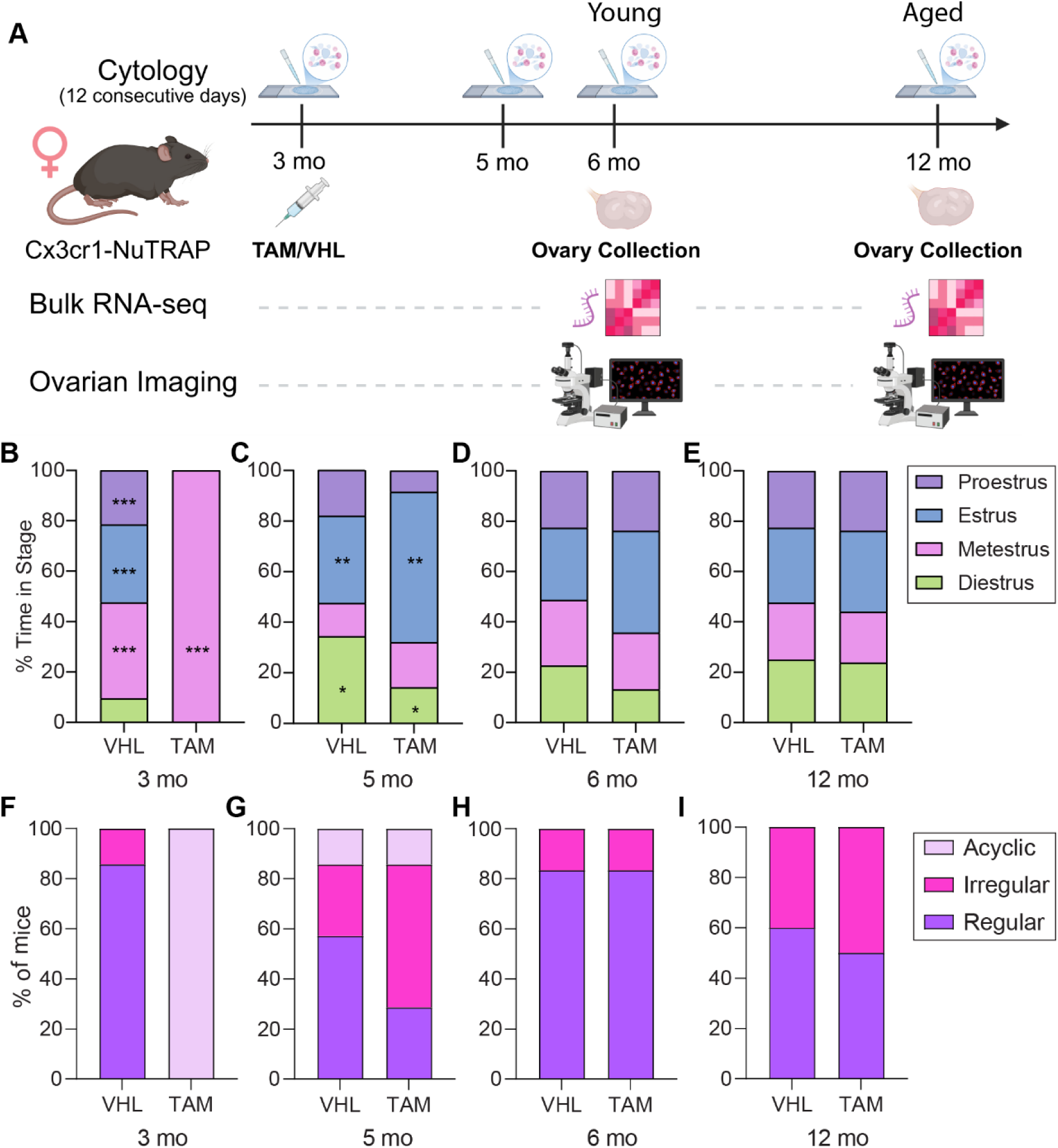
Effects of TAM administration on estrous cyclicity and ovarian outcomes in female mice. **(A)** Experimental design to test the effect of tamoxifen administration on estrous cycle and ovarian outcomes. Female Cx3cr1-NuTRAP mice were treated with Tamoxifen (TAM) or vehicle (VHL) at 3 months of age (mo). Vaginal cytology was used to assess estrous cyclicity after VHL/TAM treatment at 3, 5, 6, and 12 mo. Ovaries were collected at 6 and 12 mo for bulk RNA-sequencing and ovarian imaging analyses (n=6-7/treatment/age). **(B-E)** Distribution of time spent in each estrous stage at **(B)** 3 mo, **(C)** 5 mo, **(D)** 6 mo, **(E)** and 12 mo (2-way ANOVA, Sidak’s MTC, *p<0.05, **<0.01, ***<0.001). **(F-I)** Proportion of mice classified as having regular, irregular, or acyclic estrous cycles at **(F)** 3 mo, **(G)** 5 mo, **(H)** 6 mo, and **(I)** 12 mo.

Together, these findings indicate that TAM administration causes a marked but temporary disruption of estrous cyclicity, with recovery by approximately 3 months post-treatment and no persistent differences in cycle phase distribution through 12 mo. Thus, TAM-inducible models can likely be used in ovarian aging studies when sufficient time is allowed after TAM administration for estrous cyclicity to normalize. However, studies focused on acute ovarian responses, peri-treatment hormone-sensitive phenotypes, or outcomes measured within the first several weeks after TAM exposure should interpret results with caution and include appropriate vehicle-treated controls.

### TAM treatment does not cause persistent changes in histological features of ovarian aging

Although estrous cyclicity recovered after TAM treatment, it remained possible that TAM could have longer-lasting effects on ovarian tissue remodeling or follicle dynamics that are not reflected by cycle monitoring alone. We therefore next assessed histological features of ovarian aging, including fibrosis and accumulation of multinucleated giant cells [4] in TAM– and VHL-treated mice.

First, to determine whether TAM treatment resulted in long-term alterations to ovarian aging phenotypes, we assessed ovarian fibrosis using Picrosirius Red (PSR) staining **(Fig. 2A)**. Quantification of PSR-positive area revealed no significant effect of TAM treatment on ovarian fibrosis. However, aged ovaries exhibited significantly greater PSR staining compared to young ovaries **(Fig. 2B)**, consistent with increased age-associated collagen deposition [4]. We next assessed MNGC accumulation using Sudan Black staining **(Fig. 2C)**. Quantification of Sudan Black-positive area revealed no significant effect of TAM treatment on MNGC accumulation [4]. However, Sudan Black staining was significantly increased in aged ovaries compared to young ovaries, regardless of treatment group **(Fig. 2D)**, consistent with increased age-associated MNGC accumulation [37].

**Figure 2.**
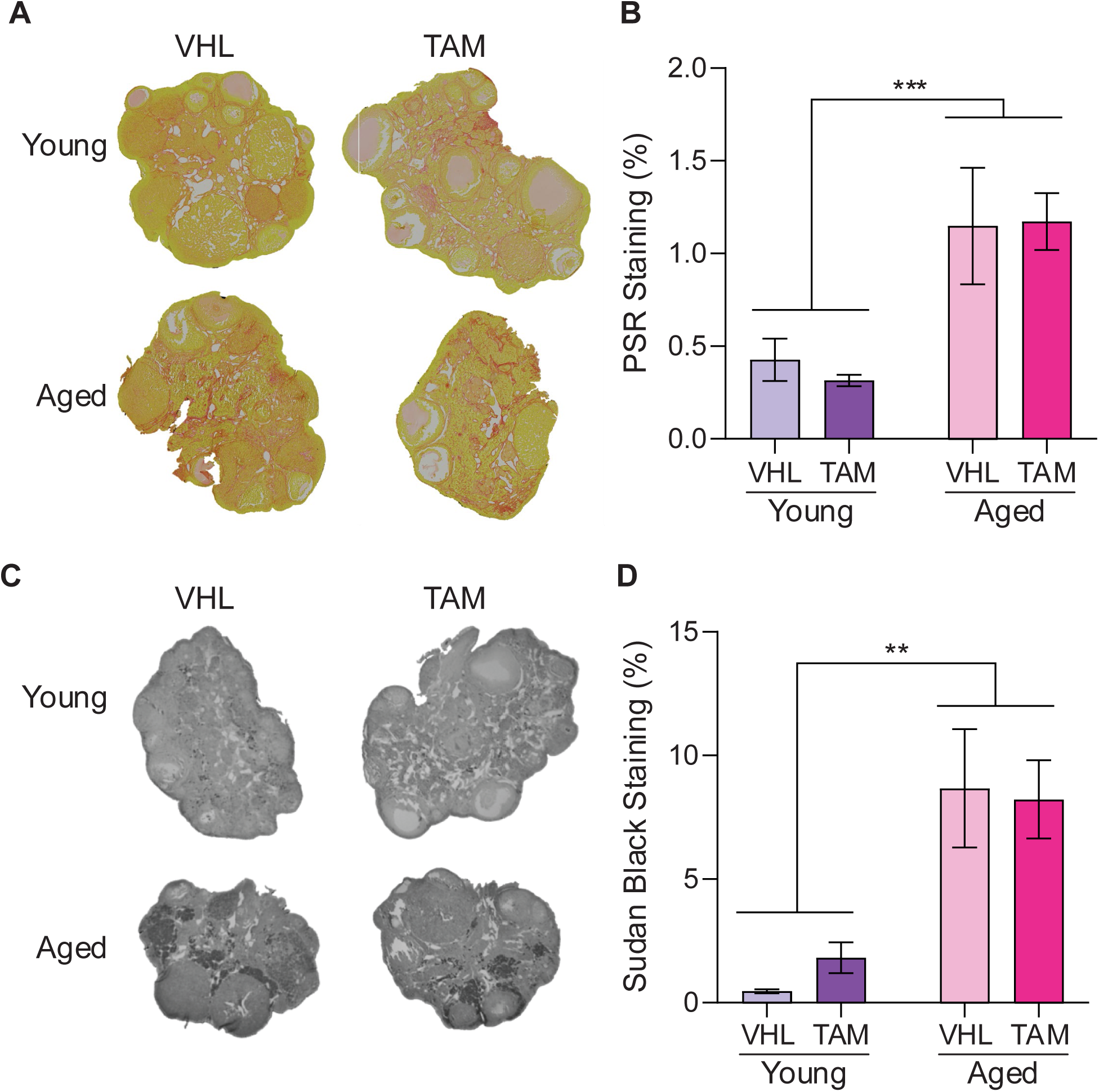
Effects of TAM administration on ovarian aging hallmarks. **(A)** Representative whole-ovary images (20X) stained with PSR to assess for fibrosis development from young and aged mice treated with VHL and TAM. **(B)** Quantification of PSR-positive area expressed as percentage of total ovarian area (2-way ANOVA, ***p< 0.001). **(C)** Representative whole-ovary images (20X) stained with Sudan Black to assess for MNGC accumulation from young and aged mice treated with VHL and TAM. **(D)** Quantification of Sudan Black-positive area expressed as percentage of total ovarian area (2-way ANOVA, **p< 0.01).

Together, these data confirm that ovarian aging is associated with increased fibrosis and MNGC accumulation regardless of TAM treatment. We found insufficient evidence to suggest that transient TAM-induced disruptions to estrous cyclicity resulted in persistent alterations to ovarian aging phenotypes

### TAM treatment is associated with limited ovarian transcriptional changes

Although estrous cyclicity recovered by 6 mo and there were no major differences in histological outcomes by treatment groups, these endpoints may not capture persistent molecular effects of TAM exposure that could influence interpretation of ovarian aging studies. Therefore, we next performed bulk RNA-seq on whole ovaries collected at 6 and 12 mo to determine whether TAM induces long-lasting transcriptional changes after apparent recovery of estrous cyclicity, with particular emphasis on pathways relevant to ovarian aging and geroscience.

We first performed differential expression analysis directly comparing TAM– and VHL-treated ovaries within each age group. This direct comparison of TAM– and VHL-treated ovaries identified few transcriptional differences. In young mice, only three genes (**Fig. 3A**) were significantly altered between treatment groups. In aged mice, three genes (**Fig. 4B**) were significantly differentially expressed between treatment groups.

**Figure 3.**
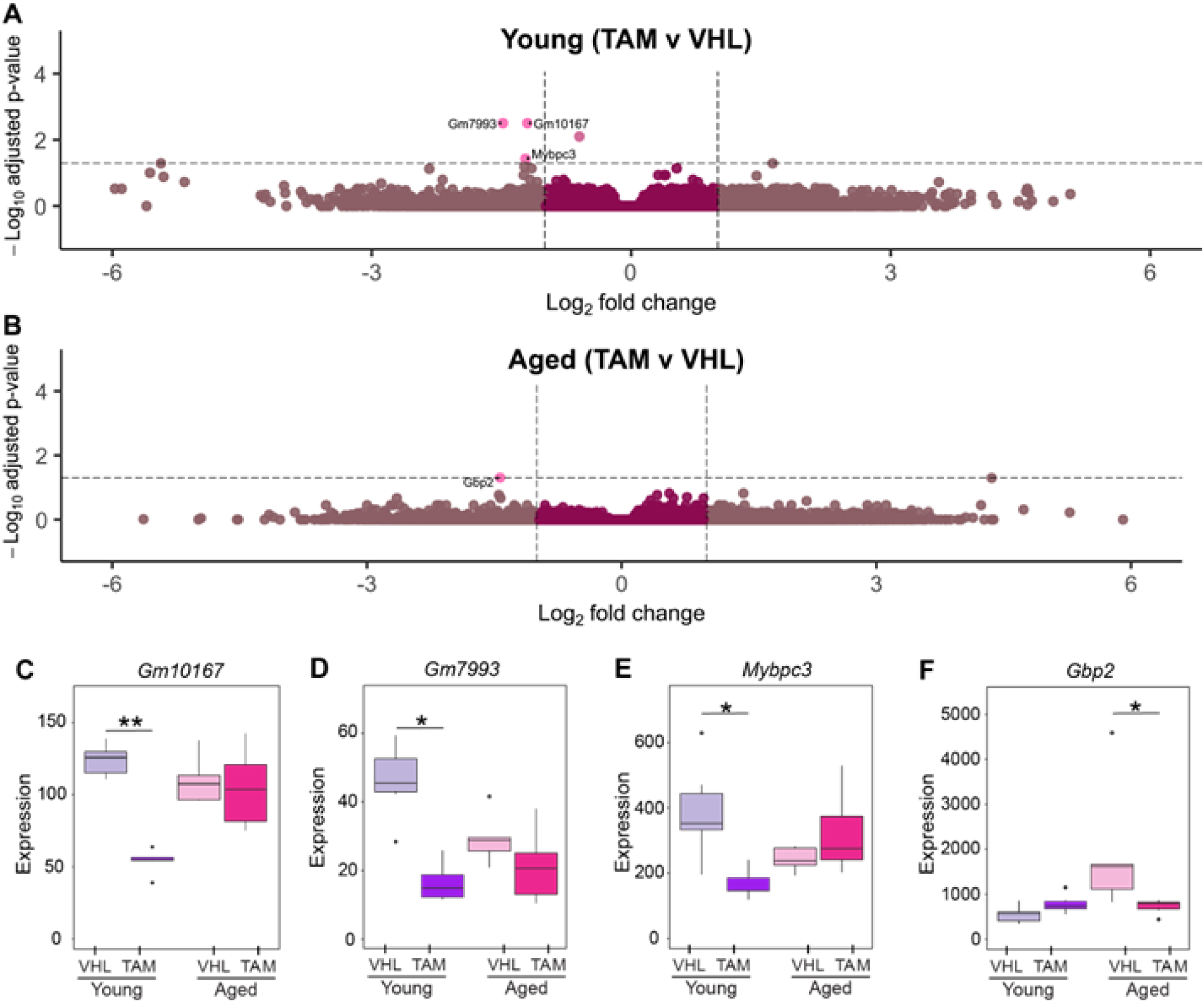
TAM treatment produces minimal alterations in ovarian transcriptional profiles. **(A)** Volcano plot showing differential gene expression between TAM– and VHL-treated young mice. **(B)** Volcano plot showing differential gene expression between TAM– and VHL-treated aged mice. **(C-F)** Gene expression of *Gm10167*, *Mybpc3*, *Gm7993*, and *Gbp2* in young and aged mice treated with VHL or TAM. Differential expression was assessed using DESeq2 with Benjamini–Hochberg-adjusted p values. DEGs were defined as adj. p< 0.05 and |log₂FC| > 1. Young VHL, n = 6; Young TAM, n = 6; Aged VHL, n = 7; Aged TAM, n = 6. * = adj. p< 0.05, ** = adj. p< 0.01.

**Figure 4.**
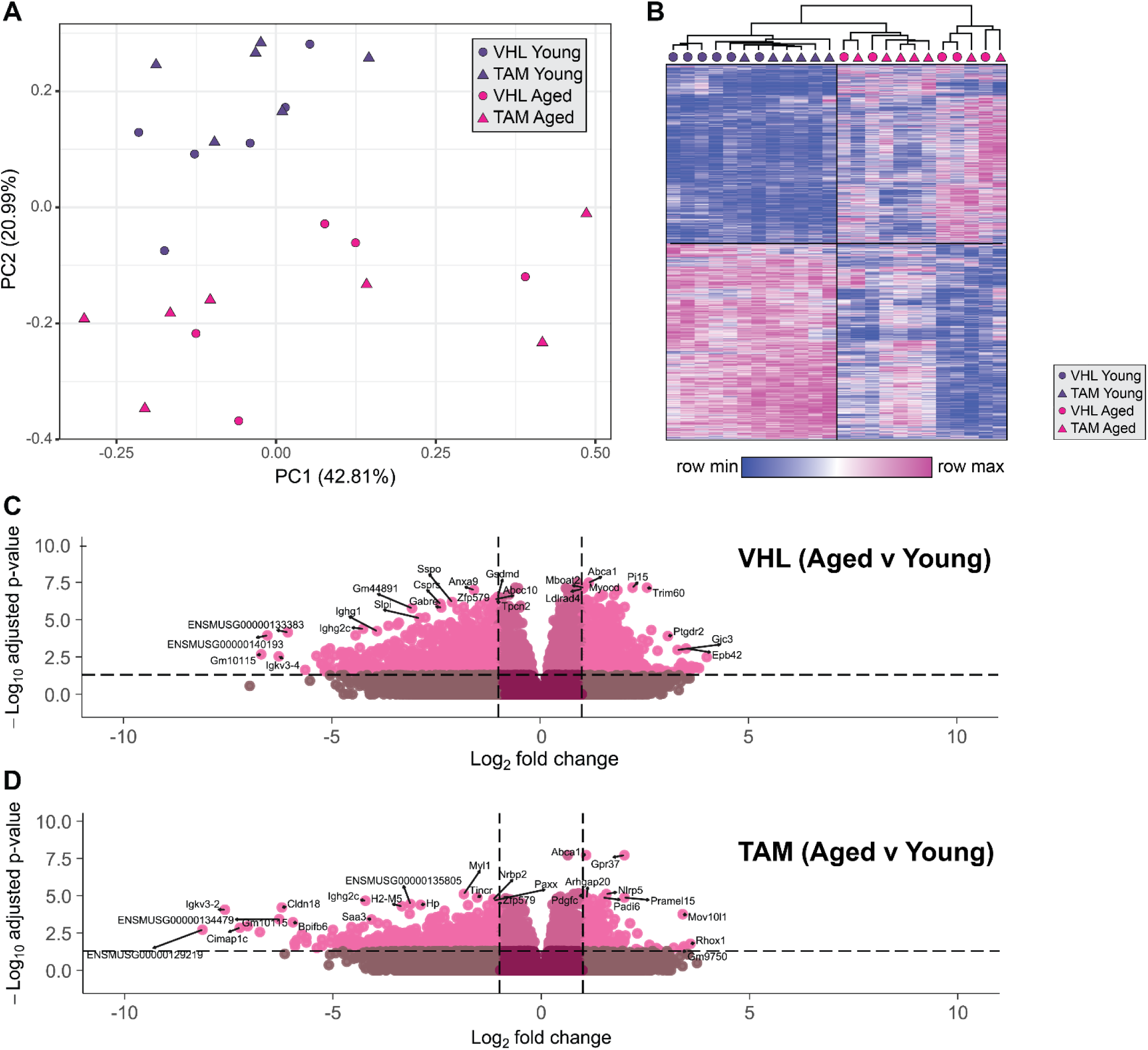
TAM treatment does not produce major long-term alterations in ovarian aging-associated transcriptional profiles. **(A)** Principal component analysis (PCA) of ovarian bulk RNA-sequencing samples from VHL– and TAM-treated young and aged Cx3cr1-NuTRAP mice. Samples clustered by age, but did not separate by treatment conditions. **(B)** Heatmap of differentially expressed genes across experimental groups. Hierarchical clustering revealed clear age-associated transcriptional differences between young and aged ovaries, while TAM and VHL groups showed broadly similar expression patterns within age groups. **(C)** Volcano plot showing differentially expressed genes in VHL-treated aged versus VHL-treated young ovaries (VHL Aged v Young). **(D)** Volcano plot showing differentially expressed genes in TAM-treated aged versus TAM-treated young ovaries (TAM Aged v Young). Differential expression was assessed using DESeq2 with Benjamini– Hochberg-adjusted P values. DEGs were defined as adj. p < 0.05 and |log₂FC| > 1. Young VHL, n = 6; Young TAM, n = 6; Aged VHL, n = 7; Aged TAM, n = 6.

Notably, the identified differentially expressed genes (DEGs) (**Fig. 3C-F**) showed limited relevance to ovarian aging. *Gm10167* and *Gm7993* (**Fig. 3C-D**) are predicted genes with poorly characterized biological functions and *Mybpc3* (**Fig. 3E**) encodes for a protein best known for its role in cardiac muscle contraction [38], its function in the ovary is unclear. Although *Gbp2* (**Fig. 3F**) is associated with interferon signaling and immune activation, it was the only DEG identified in young ovaries [39]. Overall, the small number of DEGs and the absence of clear relevant pathways suggest that TAM treatment produces little to no long-term alterations in ovarian transcriptional profiles after recovery of estrous cyclicity.

### TAM treatment does not induce persistent transcriptional changes that obscure age-related ovarian gene expression

We next compared ovarian aging trajectories between TAM– and VHL-treated mice. Principal component analysis (PCA) revealed clear separation of samples by age, whereas TAM– and VHL-treated mice were largely intermixed within each age group (**Fig. 4A**). This indicates that age was the dominant source of transcriptional variation in the ovary, with no obvious global separation attributable to prior TAM exposure. Consistent with the PCA, hierarchical clustering of DEGs showed that samples primarily clustered by age rather than treatment group (**Fig. 4B**). TAM– and VHL-treated samples were interspersed within age-defined clusters, further suggesting that prior TAM administration did not produce a strong or persistent transcriptional signature in whole ovary after recovery of estrous cyclicity.

We next compared age-associated transcriptional changes within each treatment group. Volcano plots of aged versus young ovaries in VHL– (**Fig. 4C**) and TAM-treated mice (**Fig. 4D**) showed broadly similar distributions and comparable numbers of DEGs. Together, these data suggest that ovarian aging is associated with robust transcriptional remodeling, but that prior TAM exposure does not substantially alter the overall magnitude or structure of age-related gene expression changes under these experimental conditions. There were 1487 genes that overlapped in age-associated DEGs between treatment groups with all but one of them having concordant change in direction (**Fig. 5A**). The 5,605 genes that changed in at least one group with age (**Fig. 5A**) were then correlated based on their age-related fold change in expression (**Fig. 5B**). This analysis revealed a strong correlation between age-related changes in the TAM and VHL groups, indicating that although some genes failed to meet statistical significance in both groups their overall trend in directional change was conserved after TAM treatment. These results indicate that genes altered during aging generally changed in the same direction and to a similar extent regardless of TAM treatment.

**Figure 5.**
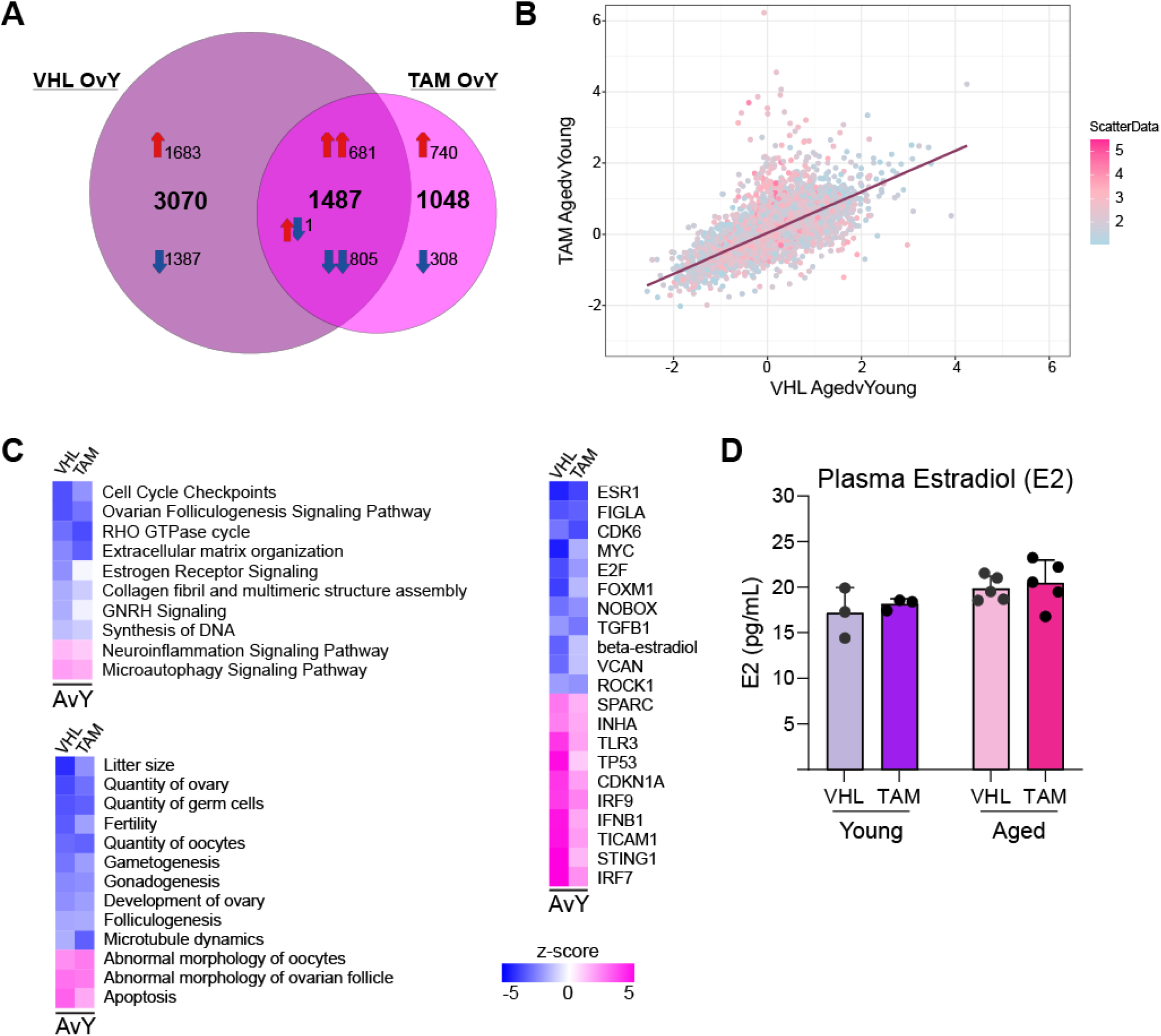
Age-associated ovarian transcriptional changes are broadly conserved between TAM– and VHL-treated mice. **(A)** Venn diagram showing overlap of differentially expressed genes (DEGs) identified in aged versus young ovaries from TAM-treated and VHL-treated mice (hypergeometric test, P < 0.001). DEGs were defined as adj. p< 0.05 and |log₂FC| > 1. **(B)** Correlation plot comparing log₂ fold-change values between VHL aged versus young and TAM aged versus young comparisons for genes significantly altered with age in at least one treatment group (Pearson r = 0.771, p< 2.2 × 10⁻¹⁶). **(C)** Heatmaps of Ingenuity Pathway Analysis (IPA) results showing predicted activation z-scores for significantly enriched diseases and biofunctions, canonical pathways, and upstream regulators associated with ovarian aging across treatment groups. **(E)** Plasma 17β-estradiol (E2) concentrations measured in young (6 mo) and aged (12 mo) mice treated with VHL or TAM. Individual animals are shown as points and bars represent mean ± SEM (2-way ANOVA).

To further examine if age-related biological processes were similarly affected between groups, we performed Ingenuity Pathway Analysis (IPA). Heatmaps of significantly enriched diseases and biofunctions, canonical pathways, and upstream regulators revealed similar patterns between TAM– and VHL-treated ovaries (**Fig. 5C**). Pathways related to ovarian aging, extracellular matrix remodeling, inflammatory signaling, folliculogenesis, and cellular stress responses showed comparable predicted activation states across treatment groups. Similarly, upstream regulators analysis identified overlapping regulatory networks associated with ovarian aging in both TAM– and VHL-treated mice. Estrogen receptor and GNRH signaling showed a modest difference in TAM-treated mice compared with VHL-treated mice. However, the difference did not correspond with broad divergence in age-associated transcriptional programs between groups and we did not detect any difference in circulating estradiol levels between the groups (**Fig. 5D**).

Together, these findings demonstrate that the molecular features of ovarian aging are broadly preserved between TAM– and VHL-treated mice, further supporting that prior TAM administration does not substantially alter long-term ovarian aging-associated transcriptional trajectories following recovery of estrous cyclicity.

### Practical recommendations for TAM-inducible Cre models in ovarian aging and cancer studies

Based on the transient disruption of estrous cyclicity observed after TAM administration and the subsequent recovery of cyclicity over time, we developed a practical framework for the use of TAM-inducible Cre models in ovarian aging studies (**Fig. 8**). These recommendations emphasize the importance of incorporating appropriate control groups, including TAM-versus VHL-treated mice and Cre-positive versus Cre-negative comparisons (when possible) to distinguish gene-specific effects from TAM-related effects. Because TAM may acutely perturb estrogen-sensitive ovarian processes, experimental designs should also standardize TAM dose, route, age at administration, and the interval between TAM exposure and endpoint collection.

**Figure 8.**
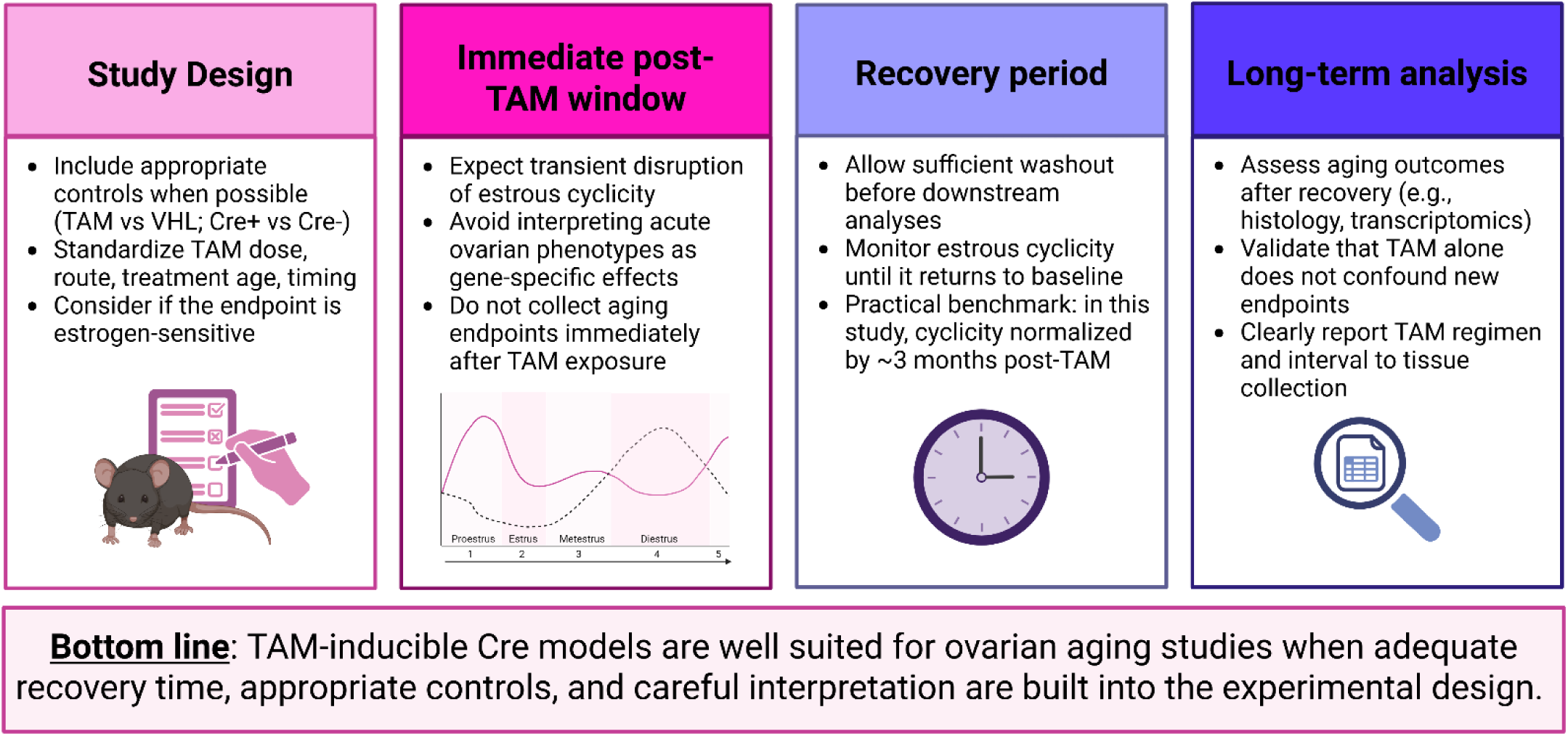
Recommendations for using TAM-inducible Cre models in ovarian aging studies. Schematic summary of key experimental recommendations for implementing TAM-inducible Cre systems in ovarian aging research. Appropriate experimental controls, including vehicle-treated and Cre-negative mice, should be incorporated and TAM administration parameters standardized. Because TAM transiently disrupts estrous cyclicity, acute ovarian phenotypes immediately following treatment should be interpreted with caution. A sufficient recovery and washout period should be included prior to downstream aging analyses, with estrous cyclicity monitored until baseline cycling resumes. Long-term aging endpoints, including histological and transcriptomic analyses, should be collected only after recovery to ensure TAM treatment itself does not confound ovarian aging-associated outcomes.

Our findings further support avoiding collection of ovarian aging endpoints during the immediate post-TAM window, when estrous cyclicity may be disrupted and acute ovarian phenotypes may reflect TAM exposure rather than the intended genetic manipulation. Instead, studies should include a sufficient recovery period before downstream analyses and, when reproductive outcomes are relevant, monitor estrous cyclicity to confirm return to baseline. In the present study, cyclicity was largely restored by approximately 3 months post-TAM treatment.

Additionally, since TAM is used clinically as a cancer therapeutic [40], proper controls should be included in ovarian cancer studies to ensure that the TAM is not influencing cancer-related endpoints.

Finally, long-term ovarian aging studies using TAM-inducible models should assess endpoints after recovery, validate that TAM alone does not confound newly examined phenotypes, and clearly report the TAM regimen and timing of tissue collection. Together, this framework supports the use of TAM-inducible Cre models in ovarian aging research when appropriate recovery time, controls, and careful interpretation are built into the experimental design.

## DISCUSSION

Tamoxifen-inducible Cre/ERT2 models are powerful tools for aging studies because they allow gene manipulation or lineage labeling to be initiated in adulthood, thereby reducing confounding effects from developmental gene alteration. This is particularly important in tissues such as the ovary, where developmental history, endocrine signaling, immune activity, and tissue remodeling all contribute to function across the reproductive lifespan. However, the same feature that makes TAM-inducible systems useful also creates an important experimental concern in ovarian studies: TAM is a selective estrogen receptor modulator and may transiently alter estrogen-sensitive ovarian processes independent of Cre-mediated recombination. Here, we evaluated whether TAM administration produces short– or long-term effects that could confound interpretation of ovarian aging outcomes.

To understand mechanisms of ovarian aging, we need models that can separate changes that happen during adulthood from changes caused earlier in development. TAM-inducible Cre/ERT2 systems are useful in this context because they allow recombination to be initiated at a defined age, enabling lineage tracing, temporal deletion or activation of genes, and cell-type-specific molecular profiling during adulthood or aging. These approaches are particularly valuable in the ovary, where many cell types contribute to both development and adult tissue remodeling. In contrast, constitutive Cre models can alter gene function throughout development, potentially disrupting tissue formation or cellular differentiation and making it difficult to determine whether later-life phenotypes reflect aging-related mechanisms, altered ovarian development, or compensatory changes that emerged earlier in life.

The Cx3cr1^Cre/ERT2^NuTRAP model was used here as a representative TAM-inducible system because similar approaches have been used in other tissues to study tissue-resident macrophages while minimizing persistent labeling of short-lived circulating monocytes [26, 41]. This distinction is especially relevant in the ovary, where both tissue-resident macrophages and monocyte-derived macrophages are present [42] but their distinct functions remain poorly defined. Because these populations can express overlapping surface markers and can be difficult to confidently resolve by lineage in single-cell transcriptomic datasets alone, inducible lineage-labeling approaches provide an important tool for investigating the specific contributions of ovarian tissue-resident macrophages independent of monocyte-derived macrophage populations.

Our results help clarify the conditions under which TAM-inducible models can be used in ovarian aging studies. Although TAM caused an acute disruption of estrous cyclicity, cycling recovered by approximately 3 months post-treatment. When ovarian outcomes were evaluated at 6 and 12 months of age, TAM-treated mice showed no major differences in ovarian fibrosis, MNGC accumulation, or whole-ovary transcriptional profiles compared with vehicle-treated controls. Instead, age was the dominant driver of both histological and transcriptional changes. These findings suggest that TAM has transient effects on ovarian function but does not appear to substantially confound the longer-term ovarian aging outcomes measured here.

An additional consideration is that TAM administration during puberty or early reproductive maturation may carry greater risk than adult induction. Puberty is a critical developmental window during which hormonal signals can exert organizational effects on multiple tissues, including the ovary and brain [43, 44]. Therefore, TAM-induced disruption of estrous cyclicity during this period could potentially alter endocrine, immune, or tissue-remodeling programs that help establish adult reproductive function. Under the treatment timeline used in this study, mice were treated at 3 months of age and evaluated after recovery at 6 and 12 months, making it difficult to assess outcomes in mice younger than 6 months without entering the acute or recovery phase after TAM exposure. In approximate human age terms, 6-month-old mice are often considered young adults (20 years), roughly analogous to peak reproductive-age humans, whereas 12-month-old mice correspond more closely to middle age (40 years) [45]. Thus, 6 months still provides an appropriate adult baseline for ovarian aging studies, although it may not capture phenotypes present during puberty or very early reproductive maturity. Investigators should therefore choose induction and collection timelines that match the biological question being asked. Future studies testing different ages at TAM administration, shorter or longer recovery intervals, and alternative routes or dosing regimens will be important for determining whether TAM-inducible models can be adapted for questions focused on pubertal development, early reproductive maturation, or acute ovarian responses. Another factor to consider in ovarian cancer studies is that TAM is used clinically as a cancer therapeutic [40]. Thus, proper controls should be included to ensure that transient TAM treatment does not impact cancer-related outcomes.

Overall, TAM-inducible models can be used effectively in ovarian aging studies, but they should not be treated as biologically neutral. Investigators should build experimental designs around the possibility of acute TAM effects by including appropriate controls, allowing sufficient recovery before endpoint collection, and validating the timing of induction for the specific ovarian phenotype being studied. This approach preserves the major advantage of inducible models while reducing the risk that transient TAM-related effects are mistaken for aging phenotypes. As interest grows in cell-specific studies of ovarian aging, TAM-inducible models offer a valuable way to target adult cell populations while minimizing developmental confounding. Our findings emphasize that these models are most appropriate when sufficient recovery time, proper controls, and endpoint-specific validation are built into the experimental design. In this context, the Cx3cr1^Cre/ERT2^NuTRAP model provides a useful strategy for studying CX3CR1-expressing cells, including efforts to distinguish tissue-resident macrophages from monocyte-derived macrophages in the aging ovary.

### Limitations of the study

A key limitation of this study is that TAM effects were evaluated using a single inducible Cre/ERT2 model, TAM dosing regimen, and age at treatment, and therefore the findings may not fully generalize to other Cre drivers, genetic backgrounds, doses, routes of administration, or experimental timelines. In addition, ovarian transcriptional analyses were performed on whole ovary, which provides an integrated tissue-level view but may miss cell-type-specific or compartment-specific effects of TAM on granulosa cells, stromal cells, immune cells, or vascular populations. Although estrous cyclicity recovered by approximately 3 months post-TAM and age remained the dominant driver of ovarian gene expression, more subtle effects on hormone levels, ovulation quality, fertility, or follicle dynamics may not be captured by the current analyses. We also did not explore the impact of TAM on ovarian cancer related outcomes. The most common and lethal form of ovarian cancer, high grade serous ovarian cancer (HGSOC), appears to arise in the fallopian tube fimbria before spreading to the ovary [46]; however, we did not evaluate the mouse oviduct, corresponding to the human fallopian tube in these studies.

### Author Contributions

S.B., E.P., and S.R.O conceived the project and designed the experiments. S.B., E.P., and H.K. performed the experiments with contributions from J.E.J.C., S.K., S.B., M.B.S., and S.R.O. S.B., S.R., and S.R.O. created figures and performed statistical analyses. S.B. and S.R.O. wrote the manuscript and all authors edited and approved the final version.

### Funding and Acknowledgements

This work was supported by the National Institutes of Health (R01 AG099844 to S.R.O. and M.B.S.) and the Global Consortium for Reproductive Longevity and Equality (GCRLE-0523 to S.R.O.; GCRLE-4501 to M.B.S.). The funders had no role in study design; data collection, analysis, or interpretation; the decision to publish; or manuscript preparation. We thank the OMRF Clinical Genomics Center, Imaging Core Facility, and Flow Cytometry and Cell Sorting Core Facility for assistance with assay optimization and instrument setup, and the OMRF Center for Biomedical Data Sciences for support with data processing and analysis. BioRender.com was used in the preparation of graphical elements included in this manuscript.

## REFERENCES

1. Camaioni, A., et al., The process of ovarian aging: it is not just about oocytes and granulosa cells. J Assist Reprod Genet, 2022. 39(4): p. 783–792.

2. Tchernof, A., et al., Menopause, central body fatness, and insulin resistance: effects of hormone-replacement therapy. Coronary Artery Disease, 1998. 9(8): p. 503–512.

3. Muka, T., et al., Association of Age at Onset of Menopause and Time Since Onset of Menopause With Cardiovascular Outcomes, Intermediate Vascular Traits, and All-Cause Mortality: A Systematic Review and Meta-analysis. JAMA Cardiology, 2016. 1(7): p. 767–776.

4. Isola, J.V.V., et al., A single-cell atlas of the aging mouse ovary. Nat Aging, 2024. 4(1): p. 145–162.

5. Zhu, Y., et al., Ovarian remodeling and aging-related chronic inflammation and fibrosis in the mammalian ovary. Journal of Ovarian Research, 2025. 18(1): p. 133.

6. Isola, J.V.V., et al., Reproductive Ageing: Inflammation, immune cells, and cellular senescence in the aging ovary. Reproduction, 2024. 168(2).

7. A K Indra1, X.W., J Brocard, J M Bornert, J H Xiao, P Chambon, D Metzger, Temporally-controlled site-specific mutagenesis in the basal layer of the epidermis: comparison of the recombinase activity of the tamoxifen-inducible Cre-ER(T) and Cre-ER(T2) recombinases<274324.pdf>. Nucleic Acids Res, 1999. Vol. 27(22).

8. Feil, S., N. Valtcheva, and R. Feil, Inducible Cre mice. Methods Mol Biol, 2009. 530: p. 343–63.

9. Robert, F., et al., Regulation of Cre Recombinase Activity by Mutated Estrogen Receptor Ligand-Binding Domains. Biochemical and Biophysical Research Communications, 1997. 237(3): p. 752–757.

10. Dutertre, M. and C.L. Smith, Molecular mechanisms of selective estrogen receptor modulator (SERM) action. J Pharmacol Exp Ther, 2000. 295(2): p. 431–7.

11. Xiao, C., J. Wang, and C. Zhang, Synthesis, Regulatory Factors, and Signaling Pathways of Estrogen in the Ovary. Reprod Sci, 2023. 30(2): p. 350–360.

12. Alvord, V.M., E.J. Kantra, and J.S. Pendergast, Estrogens and the circadian system. Semin Cell Dev Biol, 2022. 126: p. 56–65.

13. Chucair-Elliott, A.J., et al., Tamoxifen induction of Cre recombinase does not cause long-lasting or sexually divergent responses in the CNS epigenome or transcriptome: implications for the design of aging studies. Geroscience, 2019. 41(5): p. 691–708.

14. Xiao, C., J. Wang, and C. Zhang, Synthesis, Regulatory Factors, and Signaling Pathways of Estrogen in the Ovary. 2023. 30(2): p. 350–360.

15. Chucair-Elliott, A.J., et al., Inducible cell-specific mouse models for paired epigenetic and transcriptomic studies of microglia and astroglia. Commun Biol, 2020. 3(1): p. 693.

16. Quiroz, E., et al., A mouse model engineered to spatiotemporally control Cre expression in progesterone receptor positive cellsdagger. Biol Reprod, 2025. 113(1): p. 83–96.

17. Chucair-Elliott, A.J., et al., Age– and sex-divergent translatomic responses of the mouse retinal pigmented epithelium. Neurobiol Aging, 2024. 140: p. 41–59.

18. Ocanas, S.R., et al., Cell-Specific Paired Interrogation of the Mouse Ovarian Epigenome and Transcriptome. J Vis Exp, 2023(192).

19. Chassot, A.A., et al., Retinoic acid synthesis by ALDH1A proteins is dispensable for meiosis initiation in the mouse fetal ovary. Sci Adv, 2020. 6(21): p. eaaz1261.

20. Teng, K., et al., Modeling High-Grade Serous Ovarian Carcinoma Using a Combination of In Vivo Fallopian Tube Electroporation and CRISPR-Cas9-Mediated Genome Editing. Cancer Res, 2021. 81(20): p. 5147–5160.

21. Whitfield, J., et al., The estrogen receptor fusion system in mouse models: a reversible switch. Cold Spring Harb Protoc, 2015. 2015(3): p. 227–34.

22. Liu, S., et al., Construction of an aging-related risk signature in high-grade serous ovarian cancer for predicting survival outcome and immunogenicity. Medicine (Baltimore), 2023. 102(35): p. e34851.

23. Huang, S. and Q. Mo, Trends and disparities in ovarian cancer incidence and mortality in the United States, 1999 to 2023: A population-based ecological trend analysis. Medicine (Baltimore), 2026. 105(30): p. e49925.

24. Yona, S., et al., Fate mapping reveals origins and dynamics of monocytes and tissue macrophages under homeostasis. Immunity, 2013. 38(1): p. 79–91.

25. Roh, H.C., et al., Simultaneous Transcriptional and Epigenomic Profiling from Specific Cell Types within Heterogeneous Tissues In Vivo. Cell Rep, 2017. 18(4): p. 1048–1061.

26. Ocanas, S.R., et al., Minimizing the Ex Vivo Confounds of Cell-Isolation Techniques on Transcriptomic and Translatomic Profiles of Purified Microglia. eNeuro, 2022. 9(2).

27. Ekambaram, G., S.K. Sampath Kumar, and L.D. Joseph, Comparative Study on the Estimation of Estrous Cycle in Mice by Visual and Vaginal Lavage Method. J Clin Diagn Res, 2017. 11(1): p. AC05–AC07.

28. McLean, A.C., et al., Performing vaginal lavage, crystal violet staining, and vaginal cytological evaluation for mouse estrous cycle staging identification. J Vis Exp, 2012(67): p. e4389.

29. Markey, C.M., et al., Mammalian development in a changing environment: exposure to endocrine disruptors reveals the developmental plasticity of steroid-hormone target organs. Evolution & Development, 2003. 5(1): p. 67–75.

30. Ansere, V.A., et al., Cellular hallmarks of aging emerge in the ovary prior to primordial follicle depletion. Mech Ageing Dev, 2021. 194: p. 111425.

31. Briley, S.M., et al., Reproductive age-associated fibrosis in the stroma of the mammalian ovary. Reproduction, 2016. 152(3): p. 245–260.

32. Dobin, A., et al., STAR: ultrafast universal RNA-seq aligner. Bioinformatics, 2013. 29(1): p. 15–21.

33. Li, H., et al., The Sequence Alignment/Map format and SAMtools. Bioinformatics, 2009. 25(16): p. 2078–9.

34. Liao, Y., G.K. Smyth, and W. Shi, featureCounts: an efficient general purpose program for assigning sequence reads to genomic features. Bioinformatics, 2014. 30(7): p. 923–30.

35. Love, M.I., W. Huber, and S. Anders, Moderated estimation of fold change and dispersion for RNA-seq data with DESeq2. Genome Biol, 2014. 15(12): p. 550.

36. Felicio, L.S., J.F. Nelson, and C.E. Finch, Longitudinal Studies of Estrous Cyclicity in Aging C57BL/6J Mice: II. Cessation of Cyclicity and the Duration of Persistent Vaginal Cornification. Biology of Reproduction, 1984. 31(3): p. 446–453.

37. Foley, K.G., M.T. Pritchard, and F.E. Duncan, Macrophage-derived multinucleated giant cells: hallmarks of the aging ovary. Reproduction, 2021. 161(2): p. V5–V9.

38. Previs, M.J., et al., Molecular mechanics of cardiac myosin-binding protein C in native thick filaments. Science, 2012. 337(6099): p. 1215–8.

39. MacMicking, J.D., IFN-inducible GTPases and immunity to intracellular pathogens. Trends Immunol, 2004. 25(11): p. 601–9.

40. Hurteau, J.A., et al., Randomized phase III trial of tamoxifen versus thalidomide in women with biochemical-recurrent-only epithelial ovarian, fallopian tube or primary peritoneal carcinoma after a complete response to first-line platinum/taxane chemotherapy with an evaluation of serum vascular endothelial growth factor (VEGF): A Gynecologic Oncology Group Study. Gynecol Oncol, 2010. 119(3): p. 444–50.

41. Ocañas, S.R., et al., Microglial senescence contributes to female-biased neuroinflammation in the aging mouse hippocampus: implications for Alzheimer’s disease. J Neuroinflammation, 2023. 20(1): p. 188.

42. Li, N., et al., Two distinct resident macrophage populations coexist in the ovary. Front Immunol, 2022. 13: p. 1007711.

43. Sisk, C.L. and J.L. Zehr, Pubertal hormones organize the adolescent brain and behavior. Front Neuroendocrinol, 2005. 26(3-4): p. 163–74.

44. Colvin, C.W. and H. Abdullatif, Anatomy of female puberty: The clinical relevance of developmental changes in the reproductive system. Clin Anat, 2013. 26(1): p. 115–29.

45. Dutta, S. and P. Sengupta, Men and mice: Relating their ages. Life Sci, 2016. 152: p. 244–8.

46. Bergsten, T.M., J.E. Burdette, and M. Dean, Fallopian tube initiation of high grade serous ovarian cancer and ovarian metastasis: Mechanisms and therapeutic implications. Cancer Letters, 2020. 476: p. 152–160.

